# TVA-based modeling of short-term memory capacity, speed of processing and perceptual threshold in chronic stroke patients: Case-control differences, reliability, and sensitivity to cognitive training

**DOI:** 10.1101/725168

**Authors:** Geneviève Richard, Anders Petersen, Kristine M. Ulrichsen, Knut Kolskår, Dag Alnæs, Anne-Marthe Sanders, Erlend S. Dørum, Hege Ihle-Hansen, Jan Egil Nordvik, Lars T. Westlye

## Abstract

Attentional deficits following stroke are common and pervasive, and are important predictors for functional recovery. Attentional functions comprise a set of specific cognitive processes allowing to attend, filter and select among a continuous stream of stimuli. These mechanisms are fundamental for more complex cognitive functions such as learning, planning and cognitive control, all crucial for daily functioning, including social interactions. The distributed functional neuroanatomy of these processes is a likely explanation for the high prevalence of attentional impairments following stroke, and underscores the importance of a clinical implementation of computational approaches allowing for sensitive and specific modeling of attentional sub-processes. The Theory of Visual Attention (TVA) offers a theoretical, computational, neuronal and practical framework to assess the efficiency of visual selection performance and parallel processing of multiple objects. Here, using a whole-report experimental paradigm in a cross-sectional case-control comparison and in six repeated assessments over the course of a three-week intensive computerized cognitive training (CCT) intervention in chronic stroke patients (> 6 months since hospital admission, NIHSS < 7 at hospital discharge), we assessed the sensitivity and reliability of TVA parameters reflecting short-term memory capacity (*K*), processing speed (*C*) and perceptual threshold (*t*_0_). Cross-sectional group comparisons documented lower short-term memory capacity, slower processing speed and higher perceptual threshold in patients compared to age-matched healthy controls. Further, longitudinal analyses revealed high reliability of the TVA parameters in stroke patients, and higher processing speed at baseline was associated with larger cognitive benefits of the CCT. The results support the feasibility, reliability and sensitivity of TVA-based assessment of attentional functions in chronic stroke patients.

## Introduction

Attentional deficits following stroke are common, pervasive and persistent (Barker-Collo et al. 2010a), likely due to the distributed functional neuroanatomy supporting the range of attentional sub-functions. Specific functions of attention, such as the ability to sustain focus over an extended period, make relevant selections among various competing stimuli, or detect changes in perceptual scenes, are fundamental to more complex operations supporting everyday functions such as learning and social interactions, and are important predictors for functional recovery in stroke patients (Peers et al. 2018). For instance, attentional functions assessed at hospital discharge has been shown to be relevant for predicting future recovery (Hyndman et al. 2008), sustained visual and auditory attention measured two months after stroke was a strong predictor of long-term motor recovery (Robertson et al. 1997), and attentional abilities have been associated with language recovery after stroke (Geranmayeh et al. 2014).

Whereas the prevalence of attentional deficits following stroke is high, the reported estimates vary depending on assessment tool (Barker-Collo et al. 2010b; Xu et al. 2013), and is most likely underestimated when based on traditional bedside examination (Rinne et al. 2013). Given the high prevalence of attentional impairments in the acute and chronic stages of stroke and the relevance of attentional functions as a predictor of stroke recovery, there is a need to identify specific and reliable behavioral markers of attentional abilities in individual patients.

From a cognitive perspective, visual attention broadly refers to the joint set of cognitive processes that enables efficient and continuous selection and discrimination between relevant and irrelevant information from visual scenes of various degrees of complexity. The cognitive and computational sub-processes involved can be operationalized and assessed using various paradigms. Representing one of the most comprehensive and coherent accounts, the Theory of Visual Attention (TVA; Bundesen 1990) proposes a number of computational parameters that have been shown to be sensitive to cognitive aging (Espeseth et al. 2014; Habekost et al. 2013; Wiegand et al. 2018), as well as attentional impairments in several brain disorders (Habekost 2015; Habekost & Starrfelt 2009). Briefly, TVA offers a theoretical, computational and practical framework to assess an individual’s efficiency of visual selection performance and parallel visual processing of multiple objects. In TVA-based assessments, participants have to either report as many letters as possible (whole-report) or only a subset (partial-report). TVA assumes that the correctly reported letters are the winners of a race among all the letters in the visual field (later referred to as a biased competition model; Desimone & Duncan 1995).

By combining a cross-sectional case-control comparison and longitudinal assessment during the course of a computerized cognitive training (CCT) scheme (Kolskår et al. 2019) in chronic patients who suffered mild stroke (> 6 months since hospital admission, National Institute of Health Stroke Scale (NIHSS; Lyden et al. 2009) < 7 at hospital discharge), we assessed the sensitivity and reliability of TVA parameters reflecting short-term memory capacity (*K*), processing speed (*C*) and perceptual threshold (*t*_0_) based on a whole-report behavioral paradigm.

Based on the notion that TVA provides sensitive and specific measures of visual attention and that attentional deficits are common and pervasive following even relatively mild strokes, we hypothesized that (1) chronic stroke patients at the group level would show evidence of lower short-term memory capacity, slower processing speed, and higher perceptual threshold compared to healthy peers. Further, to assess the clinical sensitivity and relevance, (2) we tested for associations between clinical severity as indexed by NIHSS scores at hospital discharge and TVA parameters of visual attention in a chronic stage. In particular, we anticipated that higher NIHSS scores would be associated with poorer TVA performance. In order to assess the sensitivity to relevant etiological factors, we also tested for associations between TVA parameters at chronic stage and stroke subtype as classified using the Trial of Org 10172 in Acute Stroke Treatment (TOAST; Adams et al. 1993) classification system. Next, we hypothesized that (3) TVA parameters would constitute reliable and sensitive measures of specific attentional functions in chronic stroke patients in a longitudinal context. Lastly, assuming that specific attentional abilities are beneficial for learning and cognitive plasticity, we hypothesized that (4) higher attentional abilities as measured using TVA at baseline would be associated with larger cognitive improvement during the course of the CCT.

To test these hypotheses, we obtained cross-sectional data using a whole-report behavioral paradigm from 70 chronic stroke patients and 140 matched controls, as well as longitudinal data (6 sessions) from 54 chronic stroke patients who completed a randomized, double blind study aimed to test the utility of transcranial direct current stimulation (tDCS) in combination with CCT to improve cognitive performance following stroke (Kolskår et al. 2019; Richard et al. 2019; Ulrichsen et al. 2019).

## Materials and methods

Table 1 summarizes key demographics for the healthy controls and the patient group and Figure 1 depicts a schematic timeline of the study protocol.

**Table 1.**
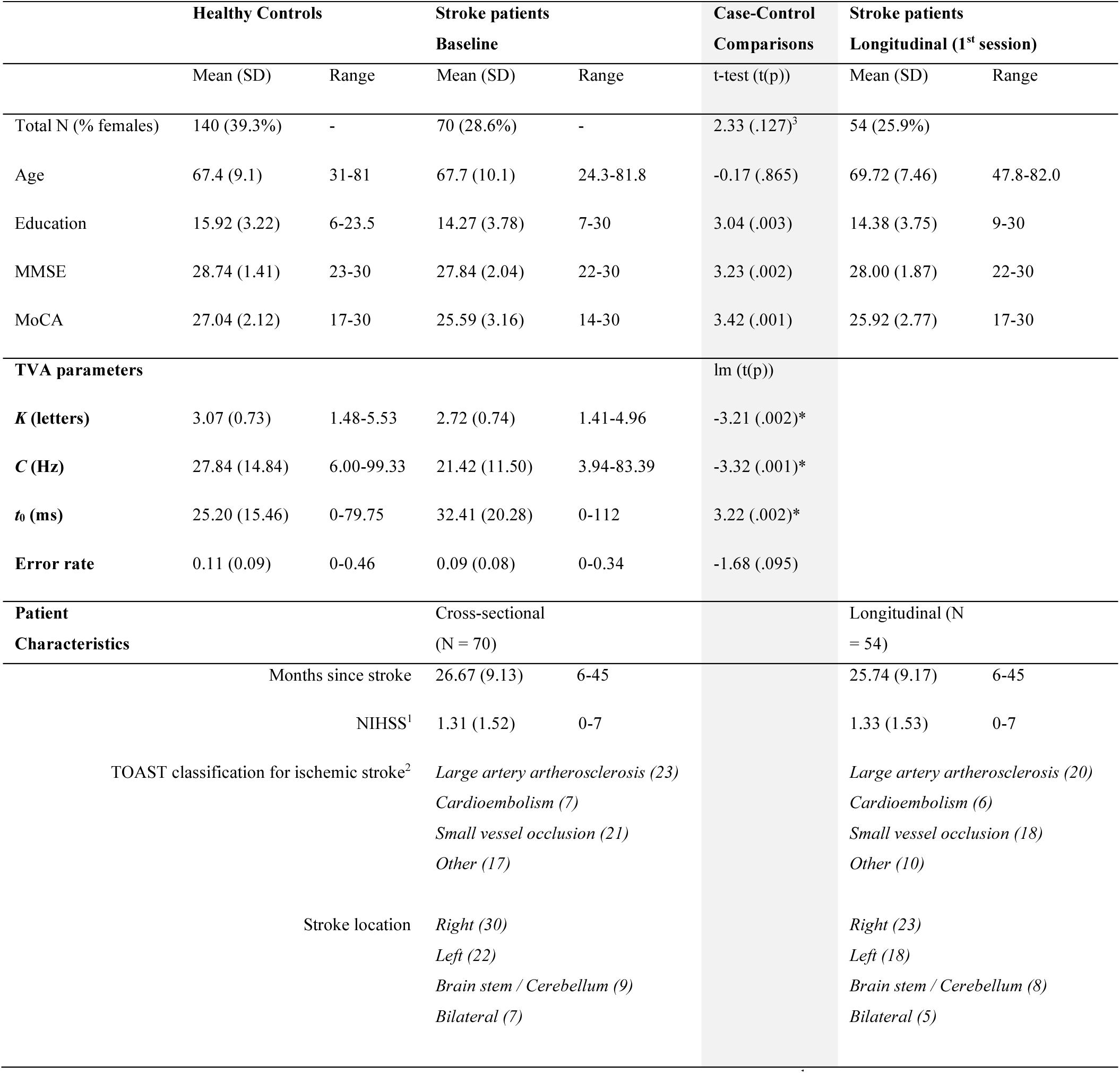
Demographics, descriptive statistics and patients sample characteristics. ^1^NIHSS score at hospital discharge. ^2^One patient had intracerebral hemorrhage. (Kolskår et al. 2019; Richard et al. 2019; Ulrichsen et al. 2019) ^3^Chi square statistics. *Significant after Bonferroni correction.

**Figure 1.**
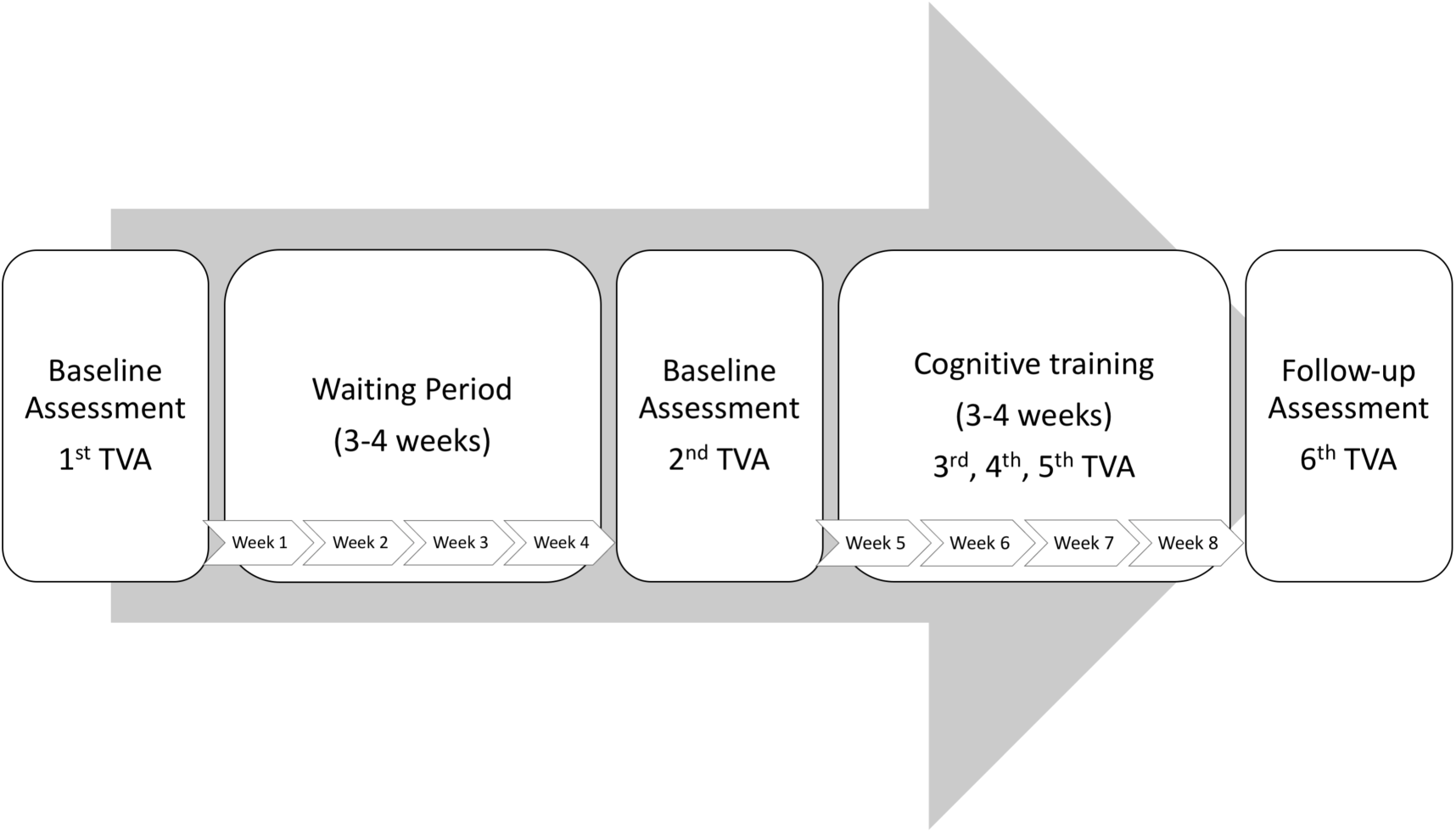
Schematic timeline of the study protocol for the stroke patients.

### Healthy controls

Healthy individuals were recruited through advertisement in newspapers, social media and word-of-mouth (Dørum et al. 2019; Richard et al. 2018). Exclusion criteria included history of stroke, dementia, or other neurologic and psychiatric diseases, alcohol- and substance abuse, medications significantly affecting the nervous system and counter indications for MRI. From a pool of 301 healthy controls who completed the behavioral paradigm and were within the relevant age range (24-81 years), we selected 140 individuals (age 31-81, mean = 67.4, SD = 9.1, 55 females) matched by age and sex using the function *matchit* with the default method *nearest* from the R package *MatchIt* (Ho et al. 2011). Here, we employed a ratio of 2:1, in which 2 healthy controls were selected for each patient.

### Patient sample

Patients admitted to the Stroke Unit at Oslo University Hospital and at Diakonhjemmet Hospital, Oslo, Norway during 2013-2016 were invited to participate in a study with the main aim to test the clinical feasibility of combining CCT and tDCS to improve cognitive function in chronic stroke patients (Kolskår et al. 2019; Richard et al. 2019; Ulrichsen et al. 2019). Stroke was defined as any form of strokes of ischemic or hemorrhagic etiology. We included patients in the chronic stage defined as a minimum of 6 months since hospital admission. Exclusion criteria included transient ischemic attacks (TIA), MRI contraindications and other neurological diseases diagnosed prior to the stroke. Seventy-two patients completed TVA-based test at the first assessment and 54 patients completed the full protocol; including three MRI brain scan sessions, three cognitive assessments, one EEG assessment, and 17 CCT sessions. Here, we excluded one patient based on incomplete baseline cognitive assessment and one due to lack of confirmed stroke resulting in the inclusion of 70 patients at baseline for the case-control comparisons (age = 24-81, mean = 67.7, SD = 10.1, 20 females), and 54 patients with longitudinal assessment (6 sessions, age = 47-82, mean = 69.7, SD = 7.5, 14 females).

The study was approved by the Regional Committee for Medical and Health Research Ethics (South-East Norway, 2014/694) and conducted in accordance with the Helsinki declaration. All participants signed an informed consent prior to enrollment and received a compensation of 500 NOK for their participation.

### CCT protocol

The computerized working memory training program (Cogmed Systems AB, Stockholm, Sweden) consisted of 25 online training sessions. In this study, we utilized 17 sessions over a period of three to four weeks, corresponding to approximately five weekly training sessions (Kolskår et al. 2019; Richard et al. 2019). On average, patients completed two training sessions combined with tDCS per week with a minimum of one day between each tDCS session. The remaining CCT sessions were performed at home. Each training session comprised eight different exercises and lasted for about 45 minutes. In total, 10 different tasks targeting verbal and visuospatial working memory were used. The difficulty level of each task was automatically adapted to the participant’s performance throughout the intervention.

### tDCS protocol

The details of the protocol have been described previously (Kolskår et al. 2019). Briefly, participants were randomly assigned to an active or a sham condition. We used a battery-driven direct current stimulator (Neuroconn DC-STIMULATOR PLUS, neuroConn GmbH, Illmenau, Germany), with the following parameters: DC current = 1 mA, total duration = 20 minutes, ramp-up = 120 seconds, fade-out = 30 seconds, and current density = 28.57 µA/cm2, rubber pads size = 5 × 7 cm. We used the factory settings for the sham condition, including a ramp-up and a fade-out period. Based on the 10-20 system for the electrode location, the anodal electrode was placed over F3 and the cathodal electrode over O2. The pads were covered with high-conductive gel (Abralyt HiCl, Falk Minow Services Herrsching, Germany) to keep the impedance threshold under < 20 kΩ and fixated with rubber bands. Side-effects were monitored following each session through self-report forms.

### TVA-based assessment

TVA-based modeling was based on data from a whole-report paradigm (Dyrholm et al. 2011; Sperling 1960), in which six red letters from a set of 20 different letters (ABDEFGHJKLMNOPRSTVXZ) were briefly presented on a circle for either 20, 40, 60, 110, and 200 ms terminated by a pattern mask or presented for 40 or 200 ms unmasked. Participants were instructed to report all the letters they were “fairly certain” of having seen (i.e., to use all available information but refrain from pure guessing). The paradigm comprised 20 practice trials and 140 test trials (i.e., 20 trials for each of the seven exposure duration conditions). TVA parameters *K*, *C*, *t*_0_ were estimated by a maximum-likelihood procedure using the LibTVA toolbox (Dyrholm et al., 2011)^1^. In addition, the error rate (i.e., the percentage of incorrect letters out of the reported letters) was calculated (hereafter referred to as an additional TVA parameter).

### Processing of Cogmed data

We used the same performance improvement scores as in previous publications (Kolskår et al. 2019; Richard et al. 2019). Briefly, we used linear modeling with performance as dependent variable and session as independent variable to quantify the changes in performance across the training period for each participant and for each trained task. Next, we performed a principal component analysis (PCA) on the performance improvement scores (zero-centered and standardized coefficients from the linear models) and used the first factor as the individual’s performance improvement to derive a common score across the trained tasks.

### Statistical analysis

Statistical analyses were performed using R version 3.3.3 (2017-03-06) (R Core Team 2017). To test our hypothesis of impaired attentional functions in stroke patients compared to healthy peers, we compared the TVA parameters between groups using linear models with each of the TVA parameters as dependent variables, group (patients and controls) as independent variable, and age and sex as covariates. To control for the number of multiple comparisons, we employed Bonferroni correction with α = 0.05/4.

To test for associations between clinical characteristics and severity at hospital discharge and TVA parameters, we used linear models with each of the TVA parameters as dependent variables and each of the clinical scores (NIHSS, TOAST classification, lesion location) as independent variables, including age and sex as covariates in all models.

To assess the reliability of the TVA parameters in a longitudinal context, we estimated the intra-class coefficient (ICC) using *ICCest* function from the *ICC* R package (Wolak et al. 2012) across the six sessions.

To test if higher attentional performance at baseline was associated with larger cognitive improvement in response to the intervention, we conducted four linear models with Cogmed performance gain as dependent variable and each of the TVA parameters as independent variable, including age and sex as covariates in all models. As a supplemental analysis, we added tDCS group (sham vs experimental) as an additional variable and tested for interactions between tDCS and each of the TVA parameters on cognitive improvement.

Lastly, we also investigated whether the experimental conditions (tDCS group) led to differential performance improvement rate on TVA using linear mixed effect models (LME) to test for associations between TVA parameters and session (time) by group (active and sham).

## Results

### Cross-sectional – Case-control

Table 1 shows the statistics for the case-control comparisons and Figure 2 depicts the associations between age and each of the TVA parameters for the stroke patients and the healthy controls. At the group level, patients performed significantly poorer than healthy controls on short-term memory capacity (*K*, *β* = -.33, std.err = 0.10, *Cohen’s d = -.45*, *t*_(206)_ = -3.21, *p*_*nom*_ = .002, *p*_*adj*_ = .008), processing speed (*C*, *β* = -6.59, std.err = 1.98, *Cohen’s d = -.46*, *t*_(206)_ = -3.32, *p*_*nom*_ = .001, *p*_*adj*_ = .004) and perceptual threshold (*t*_0_, *β* = 7.83, std.err = 2.44, *Cohen’s d = .45*, *t*_(206)_ = 3.22, *p*_*nom*_ = .002, *p*_*adj*_ = .008). Error rate did not significantly differ between groups (*β* = -.02, std.err = .01, *Cohen’s d = -.23*, *t*_(206)_ = -1.68, *p*_*nom*_ = .095, *p*_*adj*_ = .380). In addition, the analysis revealed significant main effects of age on *K* (*β* = -.02, std.err = .01, *Cohen’s d = - .55*, *t*_(206)_ = -3.91, *p*_*nom*_ = .000, *p*_*adj*_ = .000), *C* (*β* = -.34, std.err = .10, *Cohen’s d = -.47*, *t*_(206)_ = -3.39, *p*_*nom*_ = .001, *p*_*adj*_ = .004) and *t*_0_ (*β* = .41, std.err = .12, *Cohen’s d = .45*, *t*_(206)_ = 3.33, *p*_*nom*_ = .001, *p*_*adj*_ = .004), and a significant main effect of sex on *t*_0_ (*β* = - 6.68, std.err = 2.40, *Cohen’s d = -.39*, *t*_(206)_ = -2.79, *p*_*nom*_ = .006, *p*_*adj*_ = .024), indicating higher *t*_0_ in female, but no significant main effect of sex on *K* (*β* = -.14, std.err = .10, *Cohen’s d = -.18*, *t*_(206)_ = -1.35, *p*_*nom*_ = .179, *p*_*adj*_ = .716) and *C* (*β* = 2.37, std.err = 1.95, *Cohen’s d = .17*, *t*_(206)_ = 1.21, *p*_*nom*_ = .226, *p*_*adj*_ = .904).

**Figure 2.**
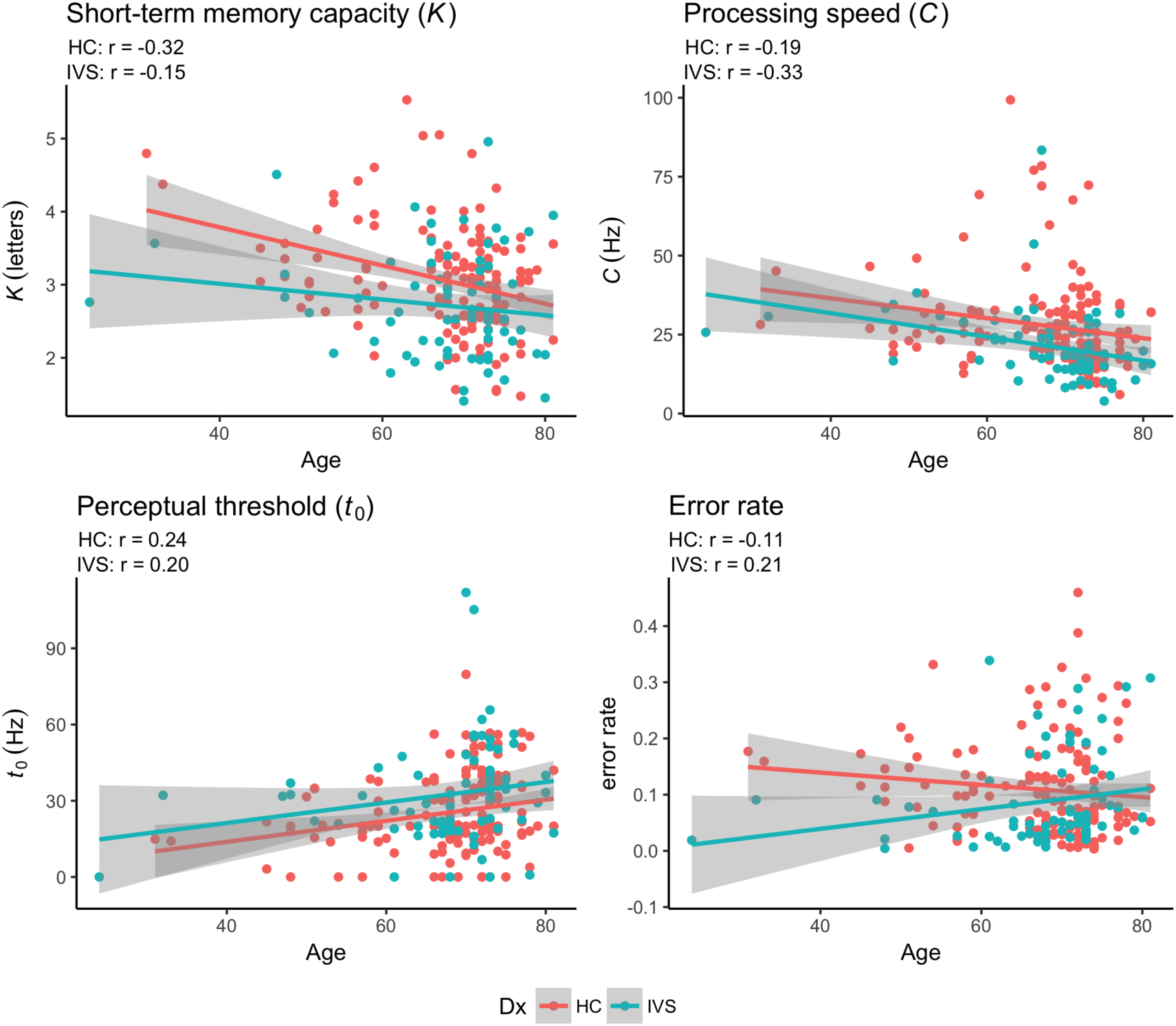
Pearson correlation between age and TVA parameters for each of the two groups independently. HC: Healthy controls. IVS: Stroke patients.

We found no significant associations between NIHSS at hospital discharge and *K* (*β* = -.066, std.err = .06, *Cohen’s d = -.27*, *t*_(59)_ = -1.037, *p*_*nom*_ = .304), *C* (*β* = .408, std.err = .98, *Cohen’s d = .11*, *t*_(59)_ = .418, *p*_*nom*_ = .678), *t*_0_ (*β* = 1.997, std.err = 1.72, *Cohen’s d = .30*, *t*_(59)_ = 1.159, *p*_*nom*_ = .251), and error rate (*β* = -.003, std.err = .01, *Cohen’s d = - .10*, *t*_(59)_ = -.376, *p*_*nom*_ = .708), and no significant main effect of TOAST classification on *K* (*F*_4,67_=1.593, *p*_*nom*_ = .188), *C* (*F*_4,67_=1.937, *p*_*nom*_ = .116), *t*_0_ (*F*_4,67_=1.188, *p*_*nom*_ = .325), and error rate (*F*_4,67_=.310, *p*_*nom*_ = .870), or of stroke location on *K* (*F*_3,68_=2.363, *p*_*nom*_ = .080), *C* (*F*_3,68_=.756, *p*_*nom*_ = .523), *t*_0_ (*F*_3,68_=.740, *p*_*nom*_ = .532), and error rate (*F*_3,68_=1.603, *p*_*nom*_ = .198).

### Longitudinal – Reliability and change over time

Figure 3 shows individual performance for each of the TVA parameters across the six timepoints for each group (sham vs tDCS) together with the global inter-class coefficients (ICCs) ranging from .58 (95% CI=.47-.69) for perceptual threshold to .80 (95% CI=.71-.86) for short-term memory capacity.

**Figure 3.**
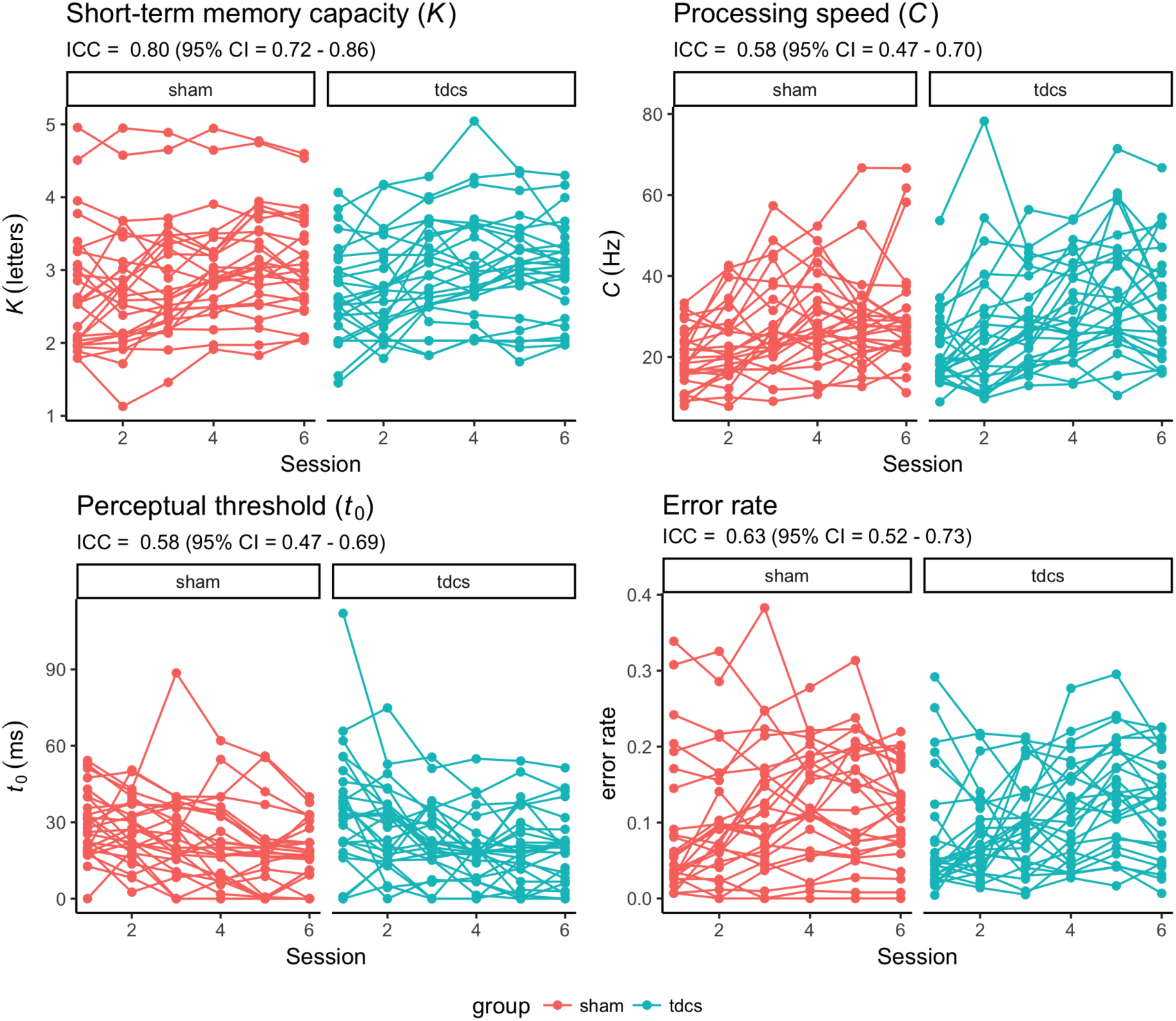
Individual performance for each of the TVA parameters across the six timepoints for each group (sham vs tDCS). Reliability of each TVA parameter is indicated by the intra-class coefficient (ICC) with 95% confidence interval (CI).

Table 2 shows summary statistics from the four linear models testing for associations between Cogmed performance gain and TVA performance at baseline. Briefly, the analysis revealed one significant association between Cogmed performance gain and processing speed (*C*) suggesting that higher processing speed at baseline was associated with larger cognitive gain during the course of the intervention. Beyond this, we found no significant effect of age or sex on performance gain. The analysis including tDCS group (sham and experimental) as an additional variable revealed no significant main effect of tDCS group, nor tDCS group by TVA performance interaction on Cogmed performance gain.

**Table 2.**
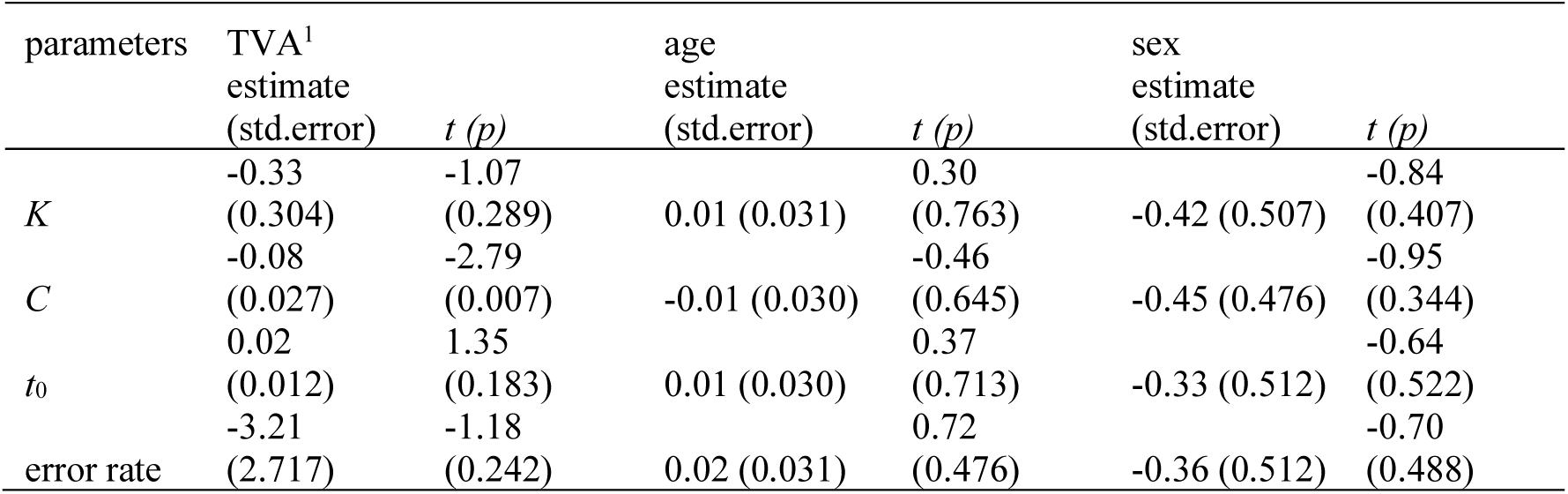
Summary statistics from the linear models testing for associations between Cogmed performance gain and TVA performance at baseline. *Significant after Bonferroni correction. ^1^Main effect of TVA parameters on Cogmed performance

Table 3 shows summary statistics from the LME testing for associations between TVA parameters and group (sham and experimental) by session interaction, including age and sex as covariates and participant as random factor. The models revealed robust main effect of session on each of the TVA parameters, suggesting improvement in performance over time. However, error rate increased over time. Beyond this, the analyses revealed a main effect of age on processing speed (*C*). We found no group by session interactions on any of the TVA parameters.

**Table 3.** Summary statistics from the linear mixed effects models testing for associations between TVA parameters and group by session, including age and sex as covariates and participant as random factor. *Significant after Bonferroni correction.

|  |  | *K* | *C* | *t*<sub>0</sub> | error rate |
| --- | --- | --- | --- | --- | --- |
| session | estimate (std.error) | 0.090 (0.014) | 2.004 (0.316) | -2.502 (0.427) | 0.006 (0.002) |
|  | *t* (*p*) | 6.62 (<.001)* | 6.34 (<.001)* | -5.86 (<.001)* | 3.18 (0.002)* |
| age | estimate (std.error) | -0.029 (0.012) | -0.666 (0.169) | 0.167 (0.248) | 0.001 (0.001) |
|  | *t* (*p*) | -2.41 (0.019) | -3.94 (<.001)* | 0.67 (0.504) | 0.96 (0.344) |
| sex | estimate (std.error) | -0.213 (0.208) | 1.967 (2.921) | -1.147 (4.286) | 0.007 (0.021) |
|  | *t* (*p*) | -1.02 (0.311) | 0.67 (0.504) | -0.27 (0.79) | 0.33 (0.744) |
| group | estimate (std.error) | 0.003 (0.194) | 1.113 (2.994) | 2.005 (4.301) | -0.024 (0.020) |
|  | *t* (*p*) | 0.02 (0.987) | 0.37 (0.712) | 0.47 (0.643) | -1.17 (0.247) |
| sess:group | estimate (std.error) | -0.018 (0.019) | 0.787 (0.446) | -0.361 (0.603) | 0.005 (0.003) |
|  | *t* (*p*) | -0.93 (0.352) | 1.76 (0.079) | -0.60 (0.550) | 1.73 (0.085) |

## Discussion

Attentional deficits are prevalent and pervasive following stroke, and are important predictors of functional and cognitive recovery. Here, we leveraged the computational framework provided by the Theory of Visual Attention (TVA) to demonstrate poorer storage capacity, lower processing speed and higher visual threshold in chronic stroke patients compared to age-matched healthy controls. Further, in a longitudinal setting, we demonstrated high reliability of the TVA parameters in stroke patients across six test sessions, and showed that higher processing speed at baseline, as indexed by the *C* parameter from TVA, was associated with larger cognitive benefits of cognitive training.

Based on the notion that TVA provides sensitive and specific measures of visual attention and that attentional deficits are common and pervasive following even relatively mild strokes, we first tested whether chronic stroke patients would show reduced performance compared to age-matched healthy controls in *K*, *C* and *t*_0_. Our results demonstrated that, at group level, patients who suffered mild stroke, defined here with a NIHSS score below seven at the hospital discharge, showed reduced performance compared to age-matched controls, suggesting that attentional deficits are present also in patients suffering from a mild stroke. This is in line with previous literature showing the TVA is sensitive to subtle deficits (Habekost & Bundesen 2003). Next, to the extent that clinical characteristics and severity of the stroke at hospital discharge are sensitive to visual attention, we tested whether higher clinical burden was associated with lower performance on TVA as measured by *K*, *C* and *t*_0_. We found no associations between NIHSS at hospital discharge, suggesting that, among patients with relatively mild strokes, the severity of the stroke is not a strong predictor of short-term memory capacity, processing speed, nor perceptual threshold as measured by TVA in a chronic stage. Further, we found no association between TOAST nor location of the stroke, and performance on TVA. Thus, neither stroke severity as measured by NIHSS at hospital discharge, nor the etiology of stroke based on TOAST or stroke location provided predictive value of attentional performance measured by TVA in a chronic stage, highlighting the complexity of the clinical etiology of long-term attentional impairments following stroke.

In line with previous findings using three test sessions in healthy individuals (Habekost et al. 2014), our results demonstrate high reliability for the *K* parameter, and fairly good reliability for *C* and *t*_0_ across the six TVA sessions in chronic stroke patients. These findings are encouraging as they support both the feasibility and reliability of computational behavioral approaches in a clinical setting, and provide support for previous cross-sectional studies implementing a similar paradigm and modeling approach.

To the extent that attentional abilities facilitate response to cognitive training, we tested whether higher attentional abilities as measured by the TVA at baseline were associated with larger improvement in response to cognitive training during the course of the intervention. Our results revealed a significant association between processing speed at baseline and performance improvement over the course of the intervention, indicating that patients with higher processing speed showed larger cognitive improvements. In contrast, visual memory capacity and perceptual threshold at baseline were not significantly associated with cognitive improvement. Thus, our results jointly demonstrate that processing speed as measured by the *C* parameter is both highly reliable, sensitive to case-control differences between stroke patients and age-matched healthy peers, and predictive for response to cognitive training in chronic stroke patients. These findings support a clinical implementation of computational cognitive approaches in general and of TVA specifically in future stroke studies.

In line with one of the major aims of the study, we also tested for effects of tDCS on cognitive improvement and interactions between attentional abilities as measured by TVA and tDCS on cognitive improvement. In addition, we also tested for tDCS by time interaction on TVA performance to assess whether the experimental conditions would result in differential improvement in TVA performance over time. Here, corroborating previous publications using the same sample (Kolskår et al. 2019; Richard et al. 2019), we found no significant associations between tDCS group (sham and experimental) and cognitive improvement, tDCS group by session interaction on TVA performance, nor tDCS group by time interaction on TVA performance, providing no significant support for the beneficial effect of tDCS.

Several methodological considerations need to be highlighted while interpreting our results. First, as previously emphasized (Richard et al. 2019), patients included in this study represent a high functioning group with relatively mild cognitive deficits and presumably better prognosis compared to patients with more severe symptoms, limiting the generalizability of our findings. It is possible that the current analysis would have revealed stronger associations between clinical variables and TVA parameters in a sample including a broader range of stroke severities and cognitive impairments. However, the labor-intensity of the current intervention and imaging study was challenging for the individual patient, preventing the study from sampling a wider clinical range of the stroke severity spectrum. Though, the value of TVA and other computational behavioral approaches may be particularly high in relatively well functioning samples where the cognitive deficits are assumed to be subtle (Ulrichsen et al. 2019). Moreover, the cross-sectional design prevents us from determining whether the group differences between patients and controls in TVA parameters primarily result from the stroke, or were present prior to the stroke. Further, the lack of control group in the longitudinal study does not allow us to separate the potential benefits from CCT on attention from the learning effect from repeated TVA assessment. Lastly, regarding the reliability estimates, although the reliability coefficients for all TVA parameters were fairly high, it is possible that the individual variability in the CCT gain could have led to underestimating ICC for the different TVA parameters. Thus, the reported values should be considered lower-bound estimates.

In conclusion, we have assessed the sensitivity and reliability of TVA parameters assessing short-term memory capacity (*K*), processing speed (*C*) and perceptual threshold (*t*_0_) derived using a whole-report behavioral paradigm in a cross-sectional case-control comparison and longitudinal assessment during the course of a CCT scheme in chronic patients who suffered mild stroke. Our results demonstrate poorer *K*, lower *C* and higher *t*_0_ in chronic stroke patients compared to age-matched healthy controls. Further, we demonstrated high reliability of the TVA parameters in stroke patients across six test sessions, and showed that higher *C* was associated with larger cognitive benefits of cognitive training. Thus, a clinically feasible implementation of TVA-based assessment offers sensitive and reliable computational parameters of short-term memory capacity, speed of processing and visual threshold in chronic stroke patients, and higher speed of processing measured at baseline was associated with stronger cognitive improvement in response to the cognitive training intervention.

## Acknowledgements and funding

This study was supported by the Norwegian ExtraFoundation for Health and Rehabilitation (2015/FO5146), the Research Council of Norway (249795, 248238), the South-Eastern Norway Regional Health Authority (2014097, 2015044, 2015073), Sunnaas Rehabilitation Hospital, and the Department of Psychology, University of Oslo.

## Footnotes

1 The model had 8 degrees of freedom (df): *K*, 5 df (the value reported is the expected *K* given a particular distribution of the probability that on a given trial, *K* = 1, 2,…, 6); *C*, 1 df; *t*_0_, 1 df; and µ (additional effective exposure duration for unmasked letters), 1 df.

